# Detection of adaptive divergence in populations of the stream mayfly *Ephemera strigata* with machine learning

**DOI:** 10.1101/424085

**Authors:** Bin Li, Sakiko Yaegashi, Thaddeus M Carvajal, Maribet Gamboa, Kozo Watanabe

## Abstract

Adaptive divergence is a key mechanism shaping the genetic variation of natural populations. A central question linking ecology with evolutionary biology concerns the role of environmental heterogeneity in determining adaptive divergence among local populations within a species. In this study, we examined adaptive the divergence among populations of the stream mayfly *Ephemera strigata* in the Natori River Basin in northeastern Japan. We used a genome scanning approach to detect candidate loci under selection and then applied a machine learning method (i.e. Random Forest) and traditional distance-based redundancy analysis (dbRDA) to examine relationships between environmental factors and adaptive divergence at non-neutral loci. We also assessed spatial autocorrelation at neutral loci to quantify the dispersal ability of *E. strigata*. Our main findings were as follows: 1) random forest shows a higher resolution than traditional statistical analysis for detecting adaptive divergence; 2) separating markers into neutral and non-neutral loci provides insights into genetic diversity, local adaptation and dispersal ability and 3) *E. strigata* shows altitudinal adaptive divergence among the populations in the Natori River Basin.

---

A central question linking ecology with evolutionary biology concerns the role of environmental heterogeneity in determining adaptive divergence among local populations within a species. Adaptive divergence in aquatic insects is usually reported to be influenced by altitudinal gradients at the river-corridor scale (Hughes et al. 2009, Keller et al. 2013, Polato et al. 2017). Altitude is often strongly related with a number of environmental factors, such as temperature and oxygen, which greatly influenced the biology of organisms (Keller and Seehausen 2012, Halbritter et al. 2015). Thermal regimes directly regulate species’ growth, development and mating behaviour, thereby setting limits on species distributions and abundances across landscapes (Li et al. 2013). Oxygen availability also restricts species’ distributions by affecting the respiratory metabolism of aquatic organisms (Rostgaard and Jacobsen 2005). Multiple studies have focused on the genetic basis of adaptive divergence in aquatic insects because of their importance in freshwater ecosystem biomonitoring. Altitudinal genetic divergence has been reported in aquatic insects including caddisflies (*Plectrocnemia conspersa* and *Polycentropus flavomaculatus* (Wilcock et al. 2007), *Stenopsyche maramorata* (Yaegashi et al. 2014), stoneflies (*Dinocras cephalotes*) (Elbrecht et al. 2014) and mayflies (Atalophlebia) (Baggiano et al. 2011). However, most of these studies were based on a given gene or a limited number of candidate genes.

The development of genome scanning approaches, such as Amplified Fragment Length Polymorphism (AFLP), allows the study of numerous anonymous markers (loci) rather than the study of a few candidate genes. Compared with neutral loci, loci influenced by directional selection (i.e. non-neutral loci) are expected to exhibit higher levels of genetic divergence (Kirk and Freeland 2011). Therefore, based on the screening of a large numbers of candidate loci (‘outlier’ loci, reviewed by Nosil et al. 2009) and the estimation of the levels of genetic divergence, statistical methods can identify loci that are under direct selection or linked to loci under selection. Selected non-neutral loci can be used to test hypotheses about the adaptive process. Neutral loci may be available for accurate tests of neutral processes, such as isolation by distance (IBD) (Oleksa et al. 2013) and gene flow patterns, thereby avoiding the confounding effects of natural selection (Kirk and Freeland 2011).

In the ordinal analysis of genome scanning, non-neutral loci are detected based on genetic variation among populations with different phenotypes or ecotypes (Bonin et al. 2006, Nosil et al. 2008, Egan et al. 2008, Galindo et al. 2009) or allopatric populations among different geographic localities (Medugorac et al. 2009, Gaggiotti et al. 2009, Renaut et al. 2011). Genome scanning can also be conducted using genetically defined populations with unknown phenotypes or ecotypes. For example, Bayesian clustering methods (Pritchard et al. 2000, Falush et al. 2003, 2007) can delineate genetic populations prior to any observable phenotypic divergence and, therefore, may provide insights into the early stages of adaptive divergence (Whiteley et al. 2011).

The determination of the link between non-neutral loci and environmental factors is one of the most difficult tasks in molecular ecology. Conventional statistical methods such as the partial Mantel test (Legendre and Fortin 2010, Watanabe et al. 2014), distance-based redundancy analysis (dbRDA) (Watanabe and Monaghan 2017) and multivariate analysis of variance (MANOVA (Mccairns and Bernatchez 2008) have been widely applied, but these methods suffer from a number of limitations. First, associating genetic variance and environmental distances can result in bias and high error rates (Legendre and Fortin 2010, Guillot and Rousset 2013, Legendre et al. 2015). In addition, the Mantel test and dbRDA are limited to testing the linear independence between genetic and environmental distances among local populations. Fulfilling the underlying assumptions of conventional statistical methods (e.g. normal distribution and homogeneity of variance) can also be very difficult (Vittinghoff et al. 2012). On account of these concerns, modern statistical techniques, such as machine learning methods, are now being developed as promising alternatives. Machine learning methods are particularly effective in finding and describing structural patterns in data and providing the values of relative importance among variables (Prasad et al. 2006, Biau and Scornet 2016).

Among the variety of machine learning methods available, Random Forest (RF) (Breiman 2001) is one of the most widely used modelling techniques to generate high-prediction accuracy and evaluate the relative importance of explanatory variables in the model (Biau and Scornet 2016). RF is an ensemble tree-based method that constructs multiple decision trees from a dataset and combines results from all the trees to create a final predictive model. In ecological studies, RF has been applied to community-level studies to predict species’ distributions and identify constrained environmental factors (Cutler 2007, Marmion et al. 2009, Evans et al. 2011). In most studies, environmental data have been used as independent variables to predict the presence or absence of species’ (dependent variables). The relative contributions of environmental variables to species distributions are quantified by their relative importance obtained from the RF model. It may therefore be possible to extend the use of RF to population genetic studies where environmental variables are used to predict the presence or absence of a haplotype or allele at outlier loci. The relative importance of each environmental variable could be considered as its influence to outlier loci, which may strongly drive adaptive divergence.

In this study, we examined adaptive divergence using AFLP markers in populations of the stream mayfly *E. strigata* from the Natori River Basin in northeastern Honshu Island, Japan (Fig.1). The primary aims of the study were to determine the extent of local adaptation at the genome level in natural populations and to quantify associations between environmental gradients and adaptive divergence. We first detected loci under selection (non-neutral loci) based on locus-specific genetic differentiation among populations. Rather than defining populations a priori using geographic or phenotypic information, we delineated populations based on the discontinuities in the AFLP variation among individuals using a hierarchical analysis of STRUCTURE (Pritchard et al. 2000, Falush et al. 2003, 2007, Vähä et al. 2007). Focusing on non-neutral loci, we then applied RF to identify environmental variables most likely to contribute to adaptive divergence and compared our results with a traditional dbRDA to examine the feasibility of the method. Finally, we examined the dispersal patterns and dispersal distance in *E. strigata* using neutral loci.

**Figure 1.**
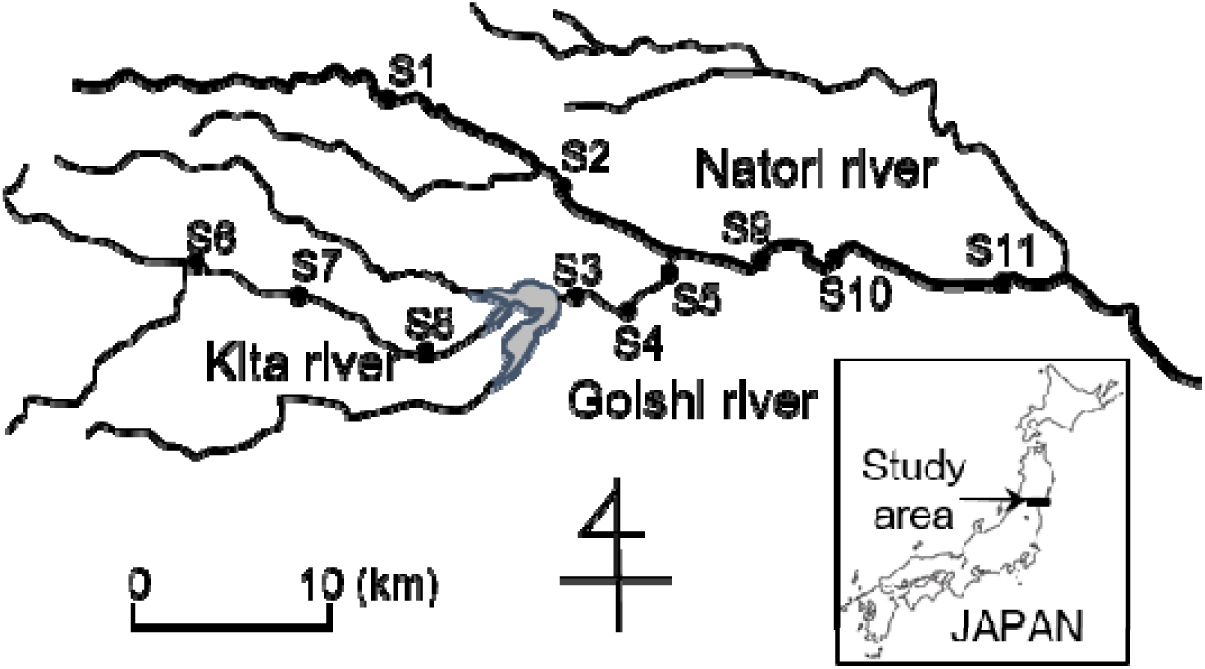
Map of 11 sampling sites for *Ephemera strigata* in the Natori River Basin in northeastern Japan.

## Methods

### Study site and sampling

*E. strigata* is a burrowing mayfly well studied in Japan and Korea (Ban and Kawai, 1986; Lee et al., 2008). In this study, sampling was carried out in the Natori River catchment in the Miyagi Prefecture in northeastern Japan (Fig. 1). Larval samples were collected at 11 sites from October 26 to November 12, 2010. At each site, we collected *E. strigata* individuals using a Surber net (30 × 30 cm quadrat with mesh size 250 μm) along 200–900 m stream reaches. All specimens were preserved in the field in 99.5% ethanol, transported to the laboratory and identified to species level under a stereomicroscope (120×) using taxonomic keys (Kawai and Tanida 2005).

We measured six geographical parameters at each site using standard ecological methods in stream surveys (Hauer and Lamberti 2007, Watanabe et al. 2008). Stream order was determined using a 1:25000 map. The width of the stream channel was measured at 10 randomly selected cross-sections using a tape measure. Longitude and latitude coordinates and altitude were recorded using a global positioning system on the river side. The riverine distance between two sites was measured on Google Maps using the ruler function.

### DNA extraction and AFLP fingerprinting

DNA from each individual was isolated from abdominal tissue by removing the digestive tract using the DNeasy 96 Blood & Tissue Kits (Qiagen). The concentration of extracted DNA was measured by Nano Drop ND-1000 spectrometer (Thermo Fisher Scientific) and diluted to 50 ng/μL. We genotyped 216 individuals from 11 sites with the AFLP method (Vos et al. 1995). The restriction step followed the protocol by Watanabe et al. (2014). The ligation step was performed by adding 1 U T4 DNA ligase (New England), 0.2 μL of 100μM MseI adapter, 0.2μM of EcoRI adapter, 2 μL T4 DNA ligase buffer (10×) (New England) and up to 20 μL dH_2_O and incubating the solution at 16°C for 12 h. The sequences of the MseI adapter and EcoRI adapters followed Reisch (2007). The adapters were manually prepared as follows: 1) mixing equal molar amounts of adapter oligomer, 2) denaturing at 95°C for 5 min and 3) incubating for 10 min at room temperature. Restricted or ligated products were then diluted at a 1:19 ratio with 0.1× TE buffer. Pre-selective amplification was performed in a mixture of 0.06 μL of 100μM MseI and EcoRI primers (Reish 2007). 15 μL of AFLP Amplification Core Mix (Applied Biosystems), 4 μL of each restricted/ligated product and up to 29 uL dH_2_O. Pre-selective polymerase chain reaction (PCR) parameters followed Reish (2007). PCR products were diluted 20 times by 0.1× TE buffer.

For selective amplifications, we employed three types of primer pairs (EcoRI-AGG & MseI-CAT, EcoRI-ACC & MseI-CAC and EcoRI-AGG & MseI-CAC) that generate the most variable patterns in 64 types of selective primer pairs using three individuals. Each EcoRI primer was modified with Beckman Dye2, 3 and 4 in 5’-end. The mixture of selective PCR was 0.1 μL of 100μM MseI and EcoRI primers, 15μL of AFLP Amplification Core Mix (Applied Biosystems) and up to 20 uL dH_2_O. We followed Reich (2007) to set PCR reaction parameters.

The selective PCR products were separated by capillary gel electrophoresis using CEQ8000 (Beckman Coulter). To adjust fluorescent intensity, each fluorescent PCR product was mixed with the following proportion EcoRI-AGG & MseI-CAT 4 μL, EcoRI-ACC & MseI-CAC 2μL and EcoRI-AGG & MseI-CAC 1μL. Peak sizes of PCR products were calculated with DNA Size Standard 600 (Beckman Coulter) using the CEQ8000 software (Beckman Coulter) with default settings.

### Hierarchal STRUCTURE analysis

We defined populations based on discontinuities in AFLP variation using the individual-based Bayesian clustering method implemented in STRUCTURE v. 2.3 (Pritchard et al. 2000, Falush et al. 2003, 2007). We performed 20 runs of 50,000 iterations with a burn-in of 10,000 for each number of assumed populations (K) ranging from 1 to 15 using the admixture model and assuming correlated allele frequencies. We used a uniform prior for alpha (the parameter representing the degree of admixture) with a maximum of 10 and set Alphapropsd to 0.05. Lambda, the parameter representing the correlation in the parental allele frequencies, was estimated in a preliminary run using K = 1. The prior *F_ST_* was set to the default value (mean = 0.01; standard deviation (SD) = 0.05).

To determine the optimal K, we computed the log-likelihood (Ln P (K)) for each K and selected K with the highest standardized second order rate of change (ΔK) of Ln P (K) (Evanno et al. 2005). Although this method helps to correctly identify K in most situations, it is known to have two limitations. First, it is useful only for the uppermost level of a hierarchical genetic structure. Second, it is unable to find the best K if K = 1 (i.e. if there is no population substructure) (Evanno et al. 2005). To address these limitations, we used a hierarchical approach for STRUCTURE analysis modified from Vähä et al. (2007), which repeats the analysis at lower hierarchical levels until no substructure can be uncovered. The advantage of our method was that we used the Wilcoxon two-sample test to control the round of repeated analysis instead of checking the pattern of individual membership. Specifically, we compared the mean value of Ln P (K) from 20 runs with optimal K (as determined using ΔK) with mean Ln P (K = 1) using the Wilcoxon two-sample test (Rosenberg et al. 2001). If Ln P (K = 1) was found to be significantly lower than Ln P (K) at the optimal K, we repeated the analysis within each of the K populations. At each hierarchical level, individuals were assigned to subpopulations based on the individual membership coefficient (Pritchard et al. 2000).

### Outlier loci detection

We used two different statistical methods to identify outlier loci. Dfdist (adapted from Fdist (Beaumont and Nichols 1996)) uses coalescent simulations to generate thousands of loci evolving under a neutral model of symmetrical islands with a mean global *F_ST_* close to the observed global *F_ST_*. Mean *F_ST_* was calculated using the default method by first excluding 30% of the highest and lowest observed values. Empirical loci with *F_ST_* values significantly greater (*p* < 0.05) than the simulated distribution (generated with 50,000 loci) were considered to be outliers. Dfdist can detect both divergent selection and balancing selection, but we focused only on divergent selection in this study.

BayeScan is a hierarchical Bayesian model-based method first described in Beaumont and Balding (2004) and modified by Foll and Gaggiotti (2008) for dominant markers (available at http://cmpg.unibe.ch/software/bayescan/). The Bayesian method is based on the concept that *F_ST_* values reflect contributions from locus-specific effects, such as selection, and population-specific effects, such as local effective size and immigration rates. The main advantage of this approach is that it allows for different demographic scenarios and different amounts of genetic drift in each population (Foll and Gaggiotti 2006, 2008). Using a reversible jump Markov Chain Monte Carlo approach, the posterior probability of each locus being subjected to selection is estimated. A locus is deemed to be influenced by selection if its *F_ST_* is significantly higher or lower than the expectation provided by the coalescent simulations.

For all subsequent analyses, non-neutral loci were defined as outlier loci detected by the Dfdist and BayeScan methods at the 95% confidence level. Neutral loci were defined as loci detected by neither Dfdist nor BayeScan at the 95% thresholds. Loci detected as outliers by only one of the two methods were not considered in the further analyses.

### Analysis of genetic diversity

*F_ST_* was calculated with ARLEQUIN v. 3.1 (Excoffier et al. 2009) using: 1) all loci, 2) only neutral loci and 3) only non-neutral loci. Global heterozygosity among all populations (*H_t_*) and mean heterozygosity within populations (*H_w_*) were estimated separately for neutral and non-neutral loci with AFLP-SURV v. 1.0 (Vekemans 2002) using the Bayesian method with a uniform prior distribution of allele frequencies (Zhivotovsky 1999). Molecular variance analysis (AMOVA) was also conducted using ARLEQUIN to provide the estimates of genetic variations among and within sampling sites. For the test of IBD, we examined the correlations of pairwise *F_ST_* with geographical distance and riverine distance (i.e. distance along the watercourse) between sites using GeneAlEx v. 6.5 (Peakall and Smouse 2012). The genetic distance between each pair of sites was quantified using mean pairwise *F_ST_* for neutral and non-neutral loci using the Bayesian-estimated allele frequencies generated by AFLP-SURV.

We conducted genetic spatial autocorrelation analysis using neutral loci for geographic distance. Eight geographic distance classes defined every 4 km (from 0–4 km to 28–32 km) were used in the analysis. Individuals within the same site were considered to be separated by a distance of 0 km. We calculated Moran’s I for each distance class using GeneAlEx, where I ranges from –1 to 1, and the positive values indicate that sites within a given distance class have similar genetic structure. We used jackknifing to estimate the 95% confidence intervals.

### Adaptive divergence modelling

We determined the environmental variables that drive adaptive divergence at non-neutral loci using the RF model (Chawla et al. 2002, Maciejewski and Stefanowski 2011, Blagus and Lusa 2013). All the six environmental variables were used to predict the band presence/absence patterns at non-neutral loci. We assigned individuals from the same site to the same environmental conditions. The dataset was imbalanced because the number of individuals with band presence was not equal to that with band absence. The individuals were thus classified in two classes (i.e. presence and absence). We solved the data imbalance problem by oversampling for the minority class using the Synthetic Minority Oversampling Technique (SMOTE) (Chawla et al. 2002). SMOTE creates synthetic minority class sample units by taking the difference between the feature vector (sample) under consideration and its nearest neighbour. It then multiplies this difference by a random number between 0 and 1 and adds it to the feature vector under consideration (Chawla et al. 2002). The process was conducted using the DMwR (Torgo 2013) and randomForest packages (Liaw and Wiener 2002) in the R programme (R Core Development Team 2015). Model performance was evaluated using the area under the receiver operating characteristic curve (AUC) (Janitza et al. 2013). The AUC value typically ranged from 0.5 (random prediction) to a maximum value of 1, which represents the perfect model theoretically. As rules of thumb, an AUC value > 0.9 indicates very good model quality, a value < 0.7 indicates poor model quality, and a value ranging from 0.7 to 0.9 indicates good model quality (Baldwin 2009).

We also conducted dbRDA as a comparative ordinal method. DbRDA was performed on the ordination solutions, rather than on the distance matrices (Legendre and Fortin 2010). In this study, pairwise genetic distances at non-neutral locus among sites were used to screen environmental factors that most closely relate to genetic divergence (Watanabe et al. 2017). The best model, comprising significant predictors, was selected using forward selection with permutation tests and an inclusion threshold of α = 0.05 using the ordistep function of the vegan package (Oksanen et al. 2015) in the R programme (R Core Development Team 2015). Significant differences were tested with the anova.cca function in the vegan package.

## Results

### Hierarchical STRUCTURE analysis

Hierarchical iterations by STRUCTURE detected significant substructure until the 4^th^ iteration beyond the initial analysis (Fig. 2). In total, 14 groups were defined for the 216 *E. strigata* individuals collected in 11 sites. Most groups were widespread over the sampling sites, whereas some groups were restricted to specific sites. For example, the members of groups 2, 3 and 8 occurred only in up- and middle-stream sites (Fig. 1: upstream sites, S1 and S6-8; middle-stream sites, S2-5).

**Figure 2.**
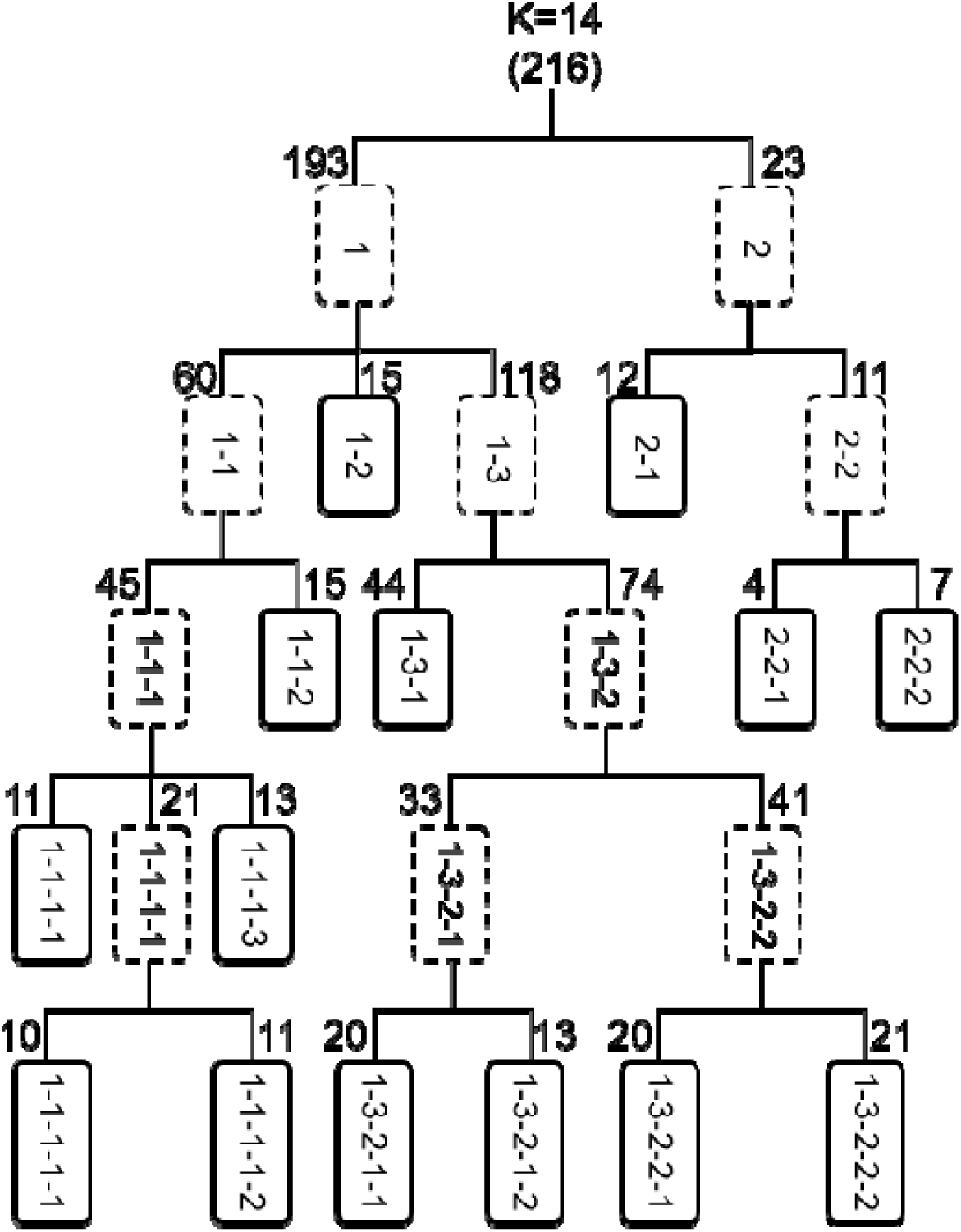
Subpopulation structure of *Ephemera strigata* as determined using STRUCTURE with hierarchical iterations. Dashed boxes indicate subpopulations and solid boxes indicate final populations. Numbers at the top of boxes indicate the number of individuals assigned to the populations. A total of 14 groups (K) were defined from 216 individuals.

### Outlier detection and genetic diversity

Using our criterion of 95% significance with both Dfdist and BayeScan, 10 non-neutral loci and 346 neutral loci were detected from the 372 polymorphic AFLP loci. Dfdist alone detected 10 outlier loci under divergent selection and 11 outlier loci under balancing selection, respectively. Outlier loci under balancing selection were not involved in this study. All the 10 outlier loci under divergent selection were consistently identified by BayeScan, which alone identified 26 outliers (Table 1). Total genetic variation (*H_t_*) was lower at neutral loci than at non-neutral loci and the same trend occurred in mean genetic variation within sites (*H_w_*; Table 2). Mean global *F_ST_* among all sites for all AFLP loci was 0.029 (*p* < 0.01; AMOVA). When measured using neutral or non-neutral loci, we found global *F_ST_* values of 0.021 (*p* < 0.01) and 0.039 (*p* < 0.01), respectively (Table 2).

**Table 1.**
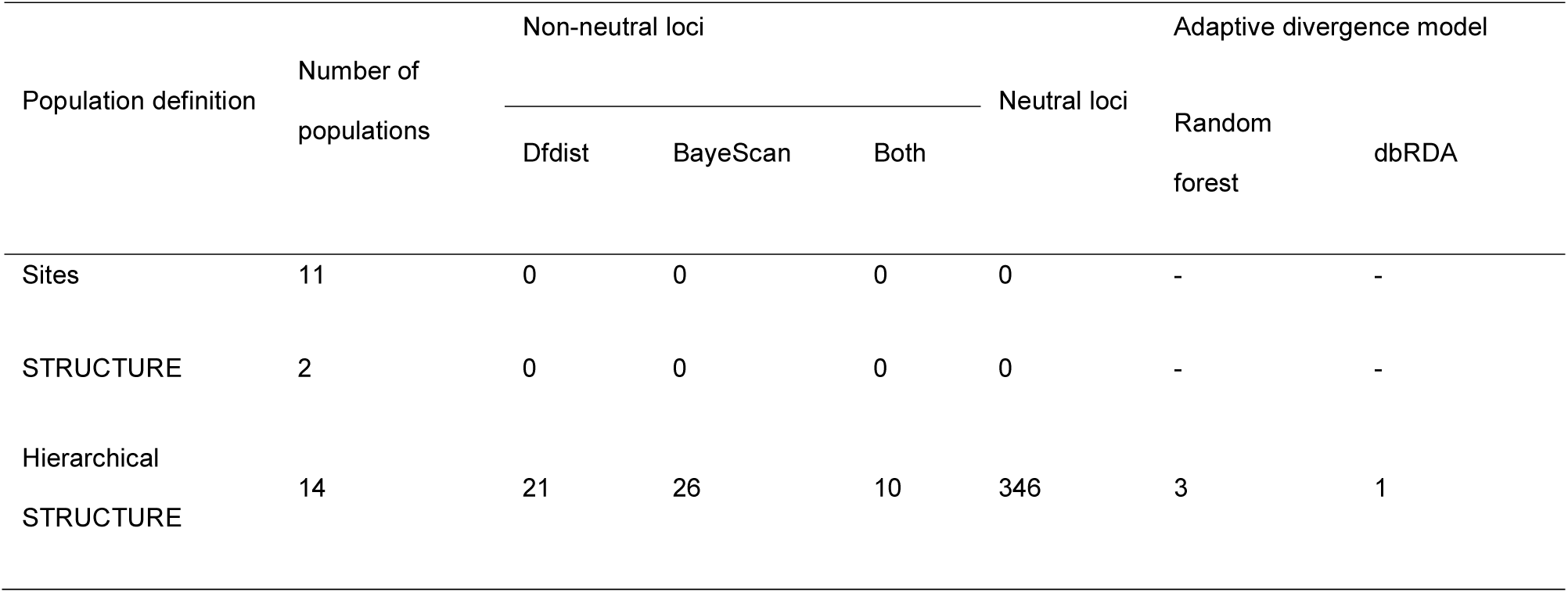
Results of outlier loci detection and model construction based on three population definitions and two adaptive divergence models. Out of the 10 non-neutral loci identified from the 14 populations delineated by the hierarchical STRUCTURE analysis, three loci (56, 89 and 254) were modelled by random forest (AUC > 0.7) and one locus (254) was modelled by dbRDA (p < 0.05).

**Table 2.** Genetic diversity and divergence measured using the following: 1) all loci, 2) only neutral loci and 3) only non-neutral loci. *H_t_* = total expected heterozygosity; *H_w_* = mean expected heterozygosity within sites; *F_ST_* = Wright’s fixation index among sites.

| | $H_t$ | $H_w$ | $F_{ST}$ |
| --- | --- | --- | --- |
| All loci | 0.1358 | 0.1357 | 0.029 |
| Neutral loci | 0.1173 | 0.1155 | 0.021 |
| Non-neutral loci | 0.4379 | 0.3523 | 0.039 |

### Detection of adaptive divergence

We separately built RF models for each of the 10 non-neutral loci. Of the 10 non-neutral loci, loci 56, 89 and 254 were well predicted (i.e. AUC > 0.7) with altitude being the most important environmental variable (Fig. 3), With dbRDA, only genetic divergence at locus 254 was significantly predicted (*p* < 0.05) (Fig. 4) with altitude explaining 54% of the genetic divergence at this locus. For the other non-neutral loci, no significant relationship with environmental factors was found with dbRDA (*p* > 0.05). IBD was not significant for both geographic (r = 0.11, *p* = 0.33) and riverine distance (r = 0.06, *p* = 0.49) (Supplementary Fig. S1). The results of the spatial autocorrelation analysis based on neutral loci showed significant positive autocorrelation coefficients at the shortest range (0–4 km; Fig. 5).

**Figure 3.**
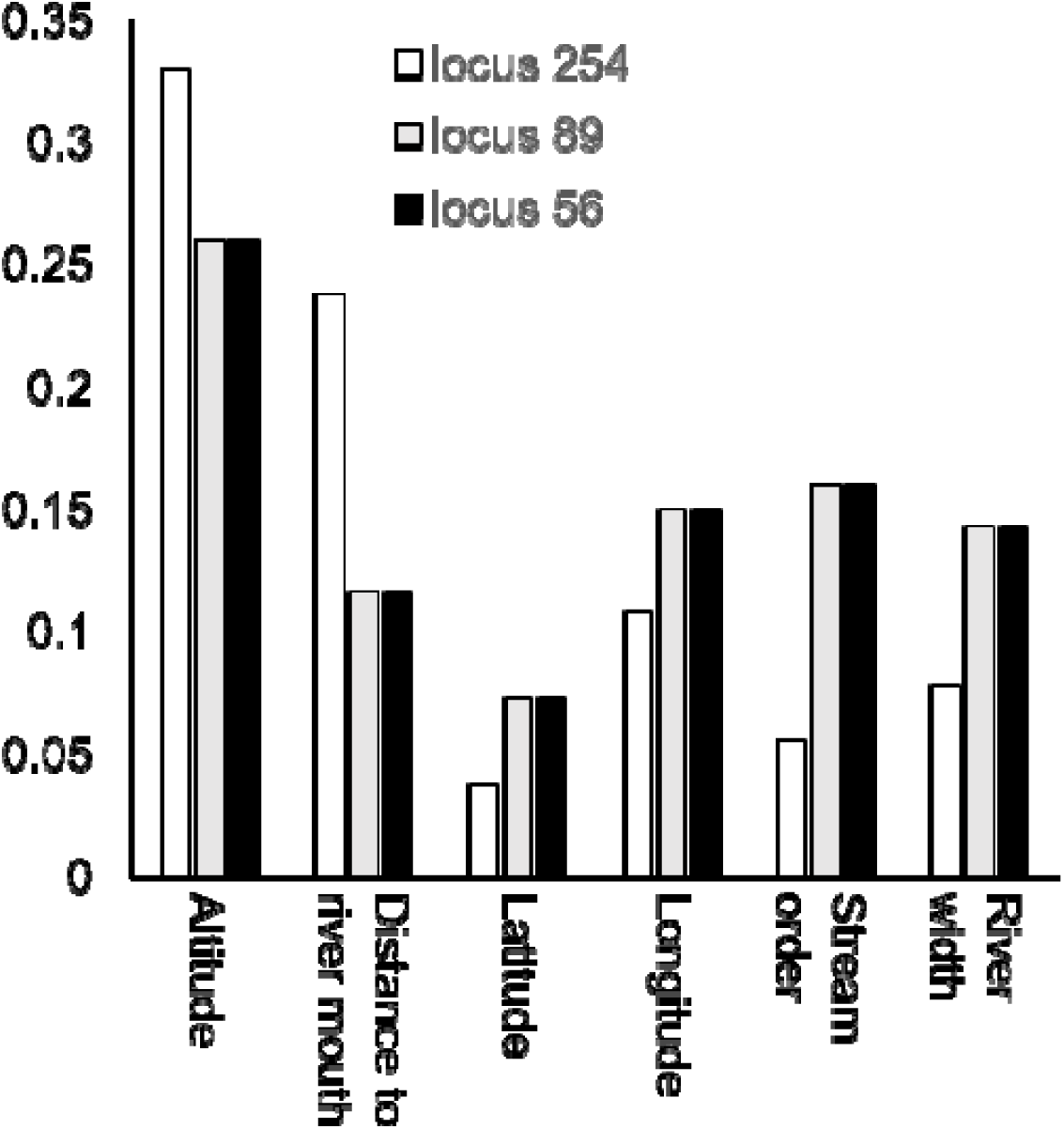
Relative importance of environmental variables based on the random forest model for three non-neutral loci (56, 89 and 254).

**Figure 4.**
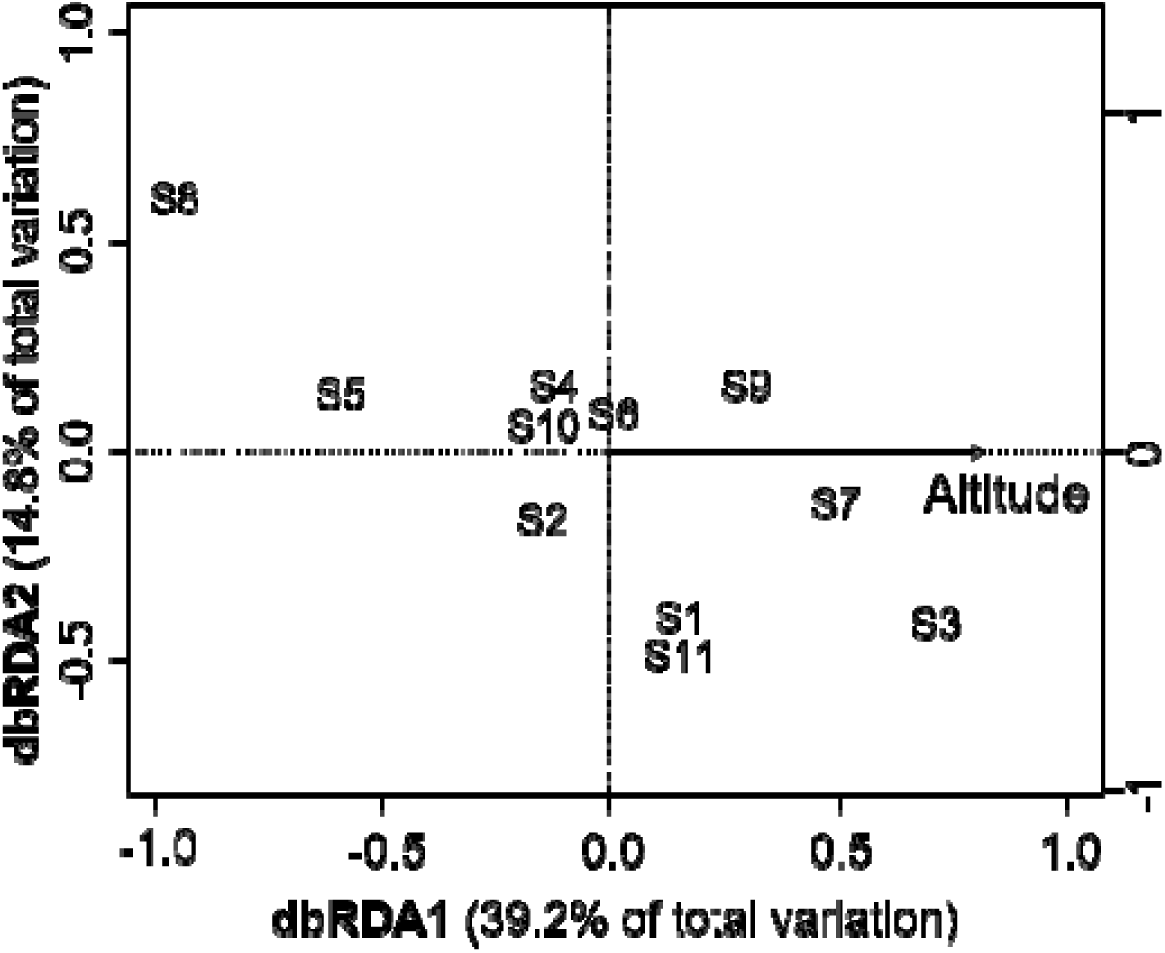
Distance-based redundancy analysis (dbRDA) describing the influence of environmental heterogeneity on genetic variation at a non-neutral locus (254).

**Figure 5.**
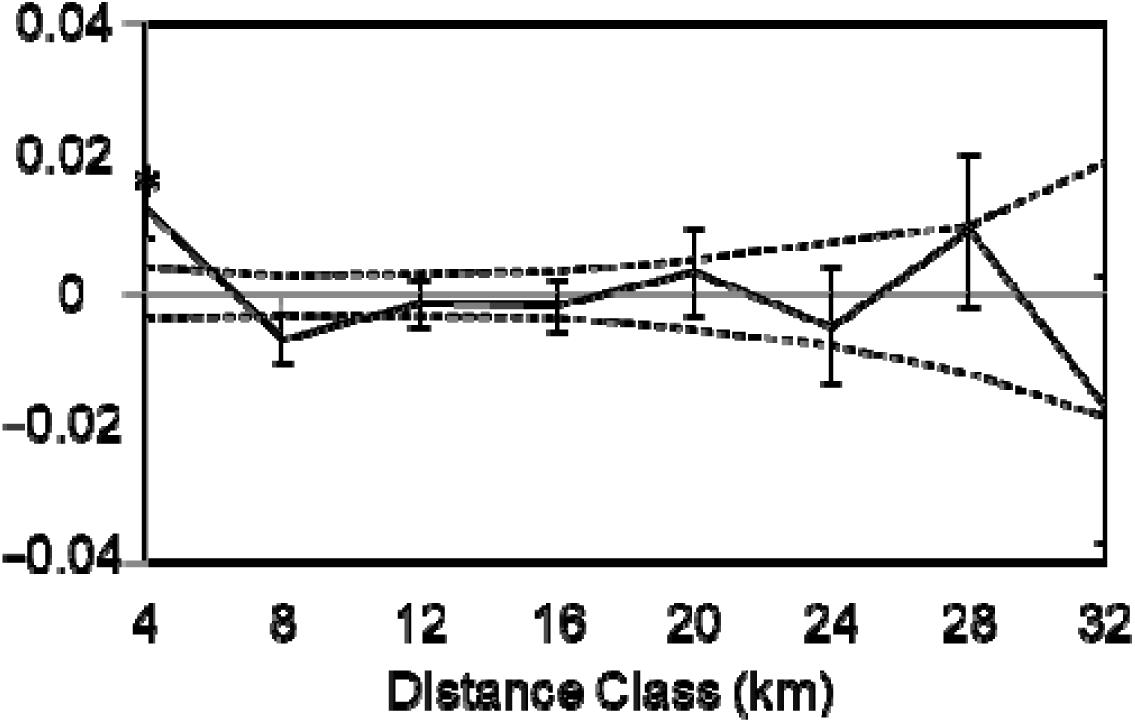
Spatial autocorrelation at 4-km distance classes based on geographic distance for neutral loci. Dashed lines indicate permutated 95% confidence intervals and error bars indicate jackknifed 95% confidence intervals. × indicates significant spatial autocorrelation (*p* < 0.05).

## Discussion

In this study, we used an RF model to examine the relationship between environmental factors and adaptive divergence at non-neutral loci in the stream mayfly *E. strigata*. Ordinal statistical tests of multiple linear regression method need assumptions that data are normally distributed with homogeneity of variance and are independent from one another (Vittinghoff et al. 2012), and this is often difficult to fulfil. The environmental factors investigated in this study did not show strong independency among variables. However, RF can overcome the limitations of regression models and accommodate pronounced nonlinearities in the exploration of gene-environment relationships in large genomic data sets (Breiman 2001, Fitzpatrick and Keller 2015, Biau and Scornet 2016). We developed RF models for each of the 10 non-neutral loci detected by both BayeScan and Dfdist. Three out of the 10 non-neutral loci (56, 89 and 254) showed good model prediction performance (AUC > 0.7), whereas the other seven could not be modelled well. This may be explained by natural selection at these seven loci being driven by environmental factors not included in our analysis (e.g. velocity and chlorophyll a) (Watanabe et al. 2014, Li et al. 2016, Brouwer et al. 2017). RF is recommended for future studies including huge numbers of environmental variables to assess their effects on adaptive divergence because RF can perform well with large numbers of variables (Genuer et al. 2010).

To compare the performance of RF with ordinal statistical analysis, we also conducted dbRDA analysis for all the 10 non-neutral loci. Only one locus (254) was well-modelled by dbRDA. This locus was one of the three loci accurately modelled by RF and the selected environmental factor (i.e. altitude) was consistent with results from RF. The low number of loci modelled in dbRDA may be because of its ability to only test linear independence^29^. The ranking of variable importance in RF relies on the principle that rearranging the values of unimportant variables should not degrade the predictive accuracy of the model (Breiman 2001). As a result, RF could reduce the influence of variable dependency on model results compared with dbRDA (Archer and Kimes 2008, Genuer et al. 2010).

To identify non-neutral loci, we used populations delineated by a hierarchal STRUCTURE analysis as an alternative to the geographic or phenotypic populations that are typically used in ordinal analysis of genome scanning. The STRUCTURE analysis successfully delineated populations with significant genetic differences, something that is difficult to achieve using visible characters such as phenotypes, ecotypes or geographic localities (Pritchard et al. 2000). The STRUCTURE analysis can delineate genetic populations among individuals prior to the occurrence of observable phenotypic divergence and may provide a means to investigate the early stages of adaptive divergence prior to phenotypic divergence in population delineation and detection of non-neutral loci (Whiteley et al. 2011).

We introduced a hierarchical approach to the STRUCTURE analysis that enabled us to look at the finer population structure (i.e. higher K) than the ordinal STRUCTURE analysis, which stops once the uppermost hierarchical level is found. The number of populations (K) is an important determinant in outlier detection (Foll and Gaggiotti 2008). We conducted outlier loci detection based on the geographical populations and uppermost hierarchical level of the STRUCTURE analysis that delineated two populations, but we could not detect any outlier loci. This clearly shows the advantages of using a hierarchical approach to STRUCTURE analysis. However, a deeper hierarchical level (e.g. the 4^th^ iteration in the hierarchy) will define a weaker structure at the risk of detecting extremely fine population substructures.

By employing a genome scanning approach, we comparatively used neutral and non-neutral loci in examining genetic diversity and genetic distance. Importantly, we found greater genetic divergence at non-neutral loci than neutral loci. This pattern is consistent with the study of three caddisflies species and one mayfly species in the same catchment system (Watanabe et al. 2014). Other studies also found similar pattern of lower levels genetic divergence in neutral DNA markers compared with morphological traits (analogues to non-neutral markers) in macroinvertebrate species such as snails (Cook 1992), spiders (Gillespie and Oxford 1998) and damselflies (Wong et al. 2003). Based on the results of Dfdist, the 10 non-neutral loci were under divergent selection rather than stabilising selection, and hence presented greater genetic divergence compared with neutral loci (Table 2).

One of the main findings of this study is that the mountain burrowing mayfly *E. strigata* presents an adaptive divergence along an altitudinal gradient. Altitude is often reported to be closely related with a number of environmental factors that influence the life cycle and development of organisms (Mórria et al. 2013, Halbritter et al. 2015). For example, altitude influences insect phenology, restricting the mating period to only a few days, thus leading to asynchronous emergence, which may act as a reproductive barrier between populations (Yaegashi et al. 2014, Watanabe et al. 2017) or as metabolism regulator (Gamboa et al. 2017). Altitude also influences air density which affects both respiration and the power required for flight. The haemoglobin gene and other genes with a potential role for adaptation to low O_2_ may show divergence between populations along an altitude gradient (Keller et al. 2013).

As opposed to non-neutral makers, neutral markers are suitable for examining neutral process occurring under the drift-migration balance. Previous population genetic studies have inferred dispersal patterns of stream insects without differentiating neutral and non-neutral loci (Miller et al. 2002, Mila et al. 2010). This may cause an overestimation of genetic drift because non-neutral loci under divergent selection will increase the estimates of genetic divergence (Kirk and Freeland 2011). Therefore, we used only neutral makers to infer dispersal patterns.

We did not find significant IBD for both geographic and riverine distances based on neutral loci, suggesting that populations are not in a genetic drift–migration equilibrium at the studied geographic scale (Supplementary Fig. S1). The results of the spatial autocorrelation analysis based on neutral loci showed significant positive autocorrelation coefficients at the shortest range (0–4 km; Fig. 5a), indicating low dispersal ability for *E. strigata*. Mayflies are generally considered to have a very low dispersal ability in mountain streams (Barber-James et al. 2007). Limited dispersal distances were also observed in stoneflies owing to their poor dispersal ability (Briers et al. 2003, 2004). In contrast, caddisflies were frequently reported to show strong dispersal ability. Yaegashi et al. (2014) reported that the caddisfly *Stenopsyche marmorata* showed pronounced dispersal ability along stream corridors up to 12 km.

In conclusion, the RF approach applied in this study performed better than the ordinal dbRDA in determining the influence of environmental factors on outlier loci under selection. Using neutral and non-neutral methods, we showed that the mountain burrowing mayfly *E. strigata* presents adaptive divergence along an altitudinal gradient. The hierarchical STRUCTURE analysis detected finer population structures and increased the power of outlier detection. A limitation of this study was that our study did not include many environmental factors, which may also be constrained factors and help to improve the model performance. In addition, sequencing the detected outlier loci would provide a deeper understanding of altitudinal adaptation in *E. strigata*.

## Acknowledgements

This research was financially supported by the Japan Society for the Promotion of Science (JSPS) (grant numbers: 16H04437, 17H01666, 16K18174). We thank K. Nagamine, S. Takahashi and Y. Kumagai for assistance with field sampling and laboratory works and T. Omura for useful suggestions. H. Harada, Tohoku University, allowed us to use their DNA sequencer and analyzing system.

## Author Contributions

B.L. analysed the data and wrote the manuscript; S.Y. conducted fieldwork, DNA extraction and AFLP experiments; T.M.C. contributed to developing the analytical methods; G.M. and K.W. edited and revised the manuscript. All authors contributed to writing the manuscript.

## Additional Information

## Supplementary Information

**Figure S1.**
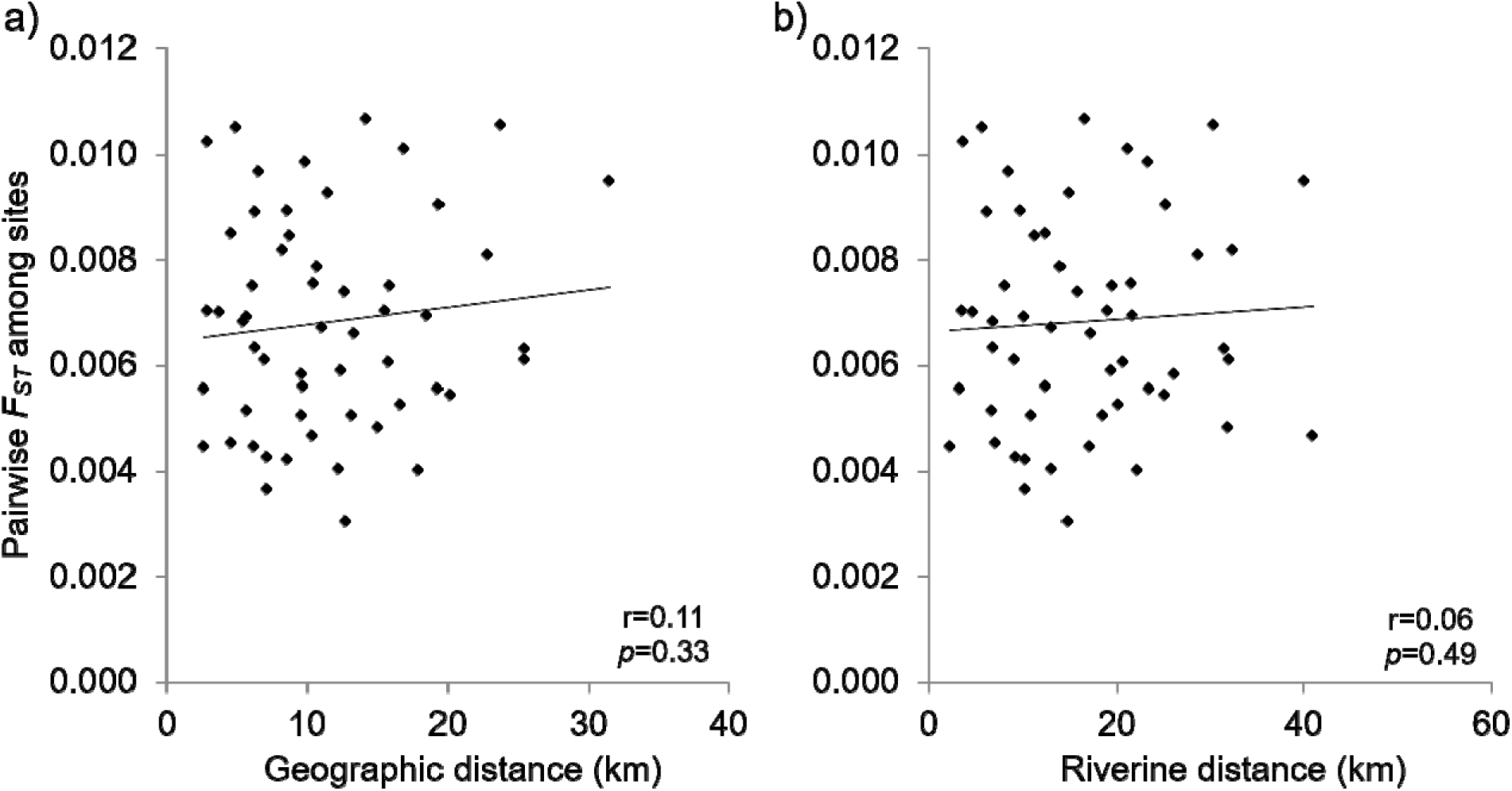
Isolation by distance calculated using geographic (a) and riverine (b) distance. Solid lines indicate correlations between Wright’s fixation index (*F_ST_*) and geographic (r = 0.11, *p* = 0.33) or riverine distance (r = 0.06, *p* = 0.49) calculated with Mantel tests.

